# Phage-centric ecological interactions in aquatic ecosystems revealed through ultra-deep metagenomics

**DOI:** 10.1101/670067

**Authors:** Vinicius S. Kavagutti, Adrian-Ştefan Andrei, Maliheh Mehrshad, Michaela M. Salcher, Rohit Ghai

## Abstract

The persistent inertia in the ability to culture environmentally abundant microbes from aquatic ecosystems represents an obstacle in disentangling the complex web of ecological interactions spun by a diverse assortment of participants (pro- and eukaryotes and their viruses). In aquatic microbial communities, the numerically most abundant actors, the viruses, remain the most elusive, and especially in freshwaters their identities and ecology remain obscure. Here, using ultra-deep metagenomic sequencing from freshwater habitats we recovered complete genomes of >2000 phages, including small “miniphages” and large “megaphages” infecting iconic prokaryotic lineages. We also show that many phages encode genes that likely enhance survival of infected microbes during strong eukaryotic grazing pressure. For instance, we describe genes that afford protection to their host from reactive oxygen species (ROS) in the environment and from the oxidative burst in protist phagolysosomes (phage-mediated ROS defense) or those that directly effect targeted killing of the predators upon ingestion of a phage-infected microbe (Trojan horse). Spatiotemporal abundance analyses of phage genomes revealed evanescence as the primary dynamic in upper water layers, where they displayed short-lived existences. In contrast, persistence was characteristic for the deeper layers where many identical phage genomes were recovered repeatedly. Phage and host abundances corresponded closely, with distinct populations displaying preferential distributions in different seasons and depths, closely mimicking overall stratification and mixis.

## Introduction

Freshwater planktonic communities are complex and dynamic, exhibiting distinct, recurrent patterns driven by both biotic and abiotic environmental factors [1]. However, in practice the accurate resolution of recurrence of individual pelagic components is challenging, and from small to large (viruses, prokaryotes, eukaryotes) our discriminative ability to quantify each participant varies greatly. Freshwaters typically contain 10^2^-10^4^ unicellular eukaryotes and 10^5^-10^7^ prokaryotes per mL, but viruses are clearly the most abundant entities, with up to 10^6^-10^8^ viruses per mL [2]. Moreover, viruses are extraordinarily diverse, and complete genomic contexts are essential to understand the nature and dynamics of this diversity. The viral collective influences microbial community ecology by increasing carbon and phosphorous transfer to microbes [3–6], modulating individual lifestyles and evolutionary histories of microbial lineages [7,8] and maintaining the diversity of the community at large [9]. Viruses in aquatic habitats are responsible for the mortality of nearly 20-40% prokaryotes every day [10], yet freshwater viruses remain largely understudied and untouched by advances in microbial culturing techniques and environmental genomics. Only a handful of isolate phage genomes are available from freshwater habitats [11–14] and only a few metagenomic studies are available [15–19]. However, the host-virus community responses to the establishment of the characteristic vertical zones in the water column of seasonally stratified water bodies, (a relatively warmer, light exposed epilimnion and a deeper and colder hypolimnion) remain uncharacterized. Even more importantly, a representative collection of complete viral genomes from freshwater has so far remained out of reach.

Here, we exploit the potential of ultra-deep metagenomic time-series sequencing to simultaneously recover phage (*Caudovirales*) and host genomic data from two common freshwater habitats (a drinking water reservoir and a humic pond). In doing so, we reconstructed 2034 complete genomes of phages infecting freshwater prokaryotes. These phage genomes are predicted to infect freshwater *Actinobacteria*, *Betaproteobacterales*, *Alphaproteobacteria*, *Bacteroidetes*, *Chloroflexi* and other phyla for which no phages have been described before. Using the abundant freshwater *Actinobacteria* and their phages as models, we show that not only do phage genome abundances in deep water bodies mirror the abundance of their hosts, they also reflect the classical patterns in thermal cycles of the water column i.e. stratification and mixis. High abundances for both phages and hosts in the epilimnion are transitory and persistence at lower abundances in the hypolimnion, the far larger niche, is the rule.

## Results and Discussion

### Metagenomic sequencing, assembly and complete phage genome recovery

We chose for our study two sites that serve as models for two distinct freshwater habitat types: meso-eutrophic Římov reservoir, a typical man-made, canyon-shaped reservoir, common to north temperate regions [20], and Jiřická pond, a shallow, humic mountain pond habitat found across the world [21]. The Římov reservoir is dimictic [22] and begins to mix at the onset of spring (March-April). It is stratified in summer when a distinct, warm epilimnion develops (May-October) above a colder hypolimnion. At the onset of winter, the colder waters sink, and the reservoir mixes again when water temperature throughout the water column drops to ca. 4ºC (Supplementary Figure S1). It is ice-covered during winter for at least two months (See methods for more site details).

We generated metagenomic time-series from both sites, producing 18 metagenomes from Římov (both epi- and hypolimnion) and 5 from Jiřická (12.97 billion reads, ca. 1.9 Tb, Supplementary Table S1). While most samples we sequenced were ca. 54 Gb in size (ranging from 190-482 million reads, average 368 million reads), two Římov samples (epi and hypolimnion) we sequenced ca. 380 Gb each (2.5 billion reads each). An overview of the microbial community using 16S rRNA abundances for both sites is shown in Supplementary Figure S2.

We also collected an additional 149 publicly available freshwater metagenomes (Supplementary Table S1, total of 4.04 billion reads, 1.09 Tb data) to search for complete phage genomes. All datasets were assembled independently (no co-assembly). In total, we analyzed ca. 3 Tb of metagenomic sequences from freshwater (ca. 17 billion reads).

The number of complete phage genomes recovered from any sample increased with sequencing depth, but with diminishing returns (Figure 1a), with genome recovery maintaining linearity up to 100 Gb (ca. 1 phage genome for every additional Gb) before tapering off at a maximum of 160 genomes from 400 Gb sequence data. While a total of 1677 genomes were assembled from the sequence data generated from the two study sites, (Římov and Jiřická), 357 genomes were recovered from all other available freshwater metagenomes. This suggests that the potential of ultra-deep sequencing to recover far more phage genomes has not yet been fully realized. We also recovered a number of metagenome-assembled genomes (MAGs) from the Římov metagenome time-series dataset (see below). We denominate this entire collection of genomes as the Uncultured Freshwater Organisms (UFO) dataset, where the UFOv subset refers to viruses and the UFOp subset to prokaryotic genomes.

**Figure 1.**
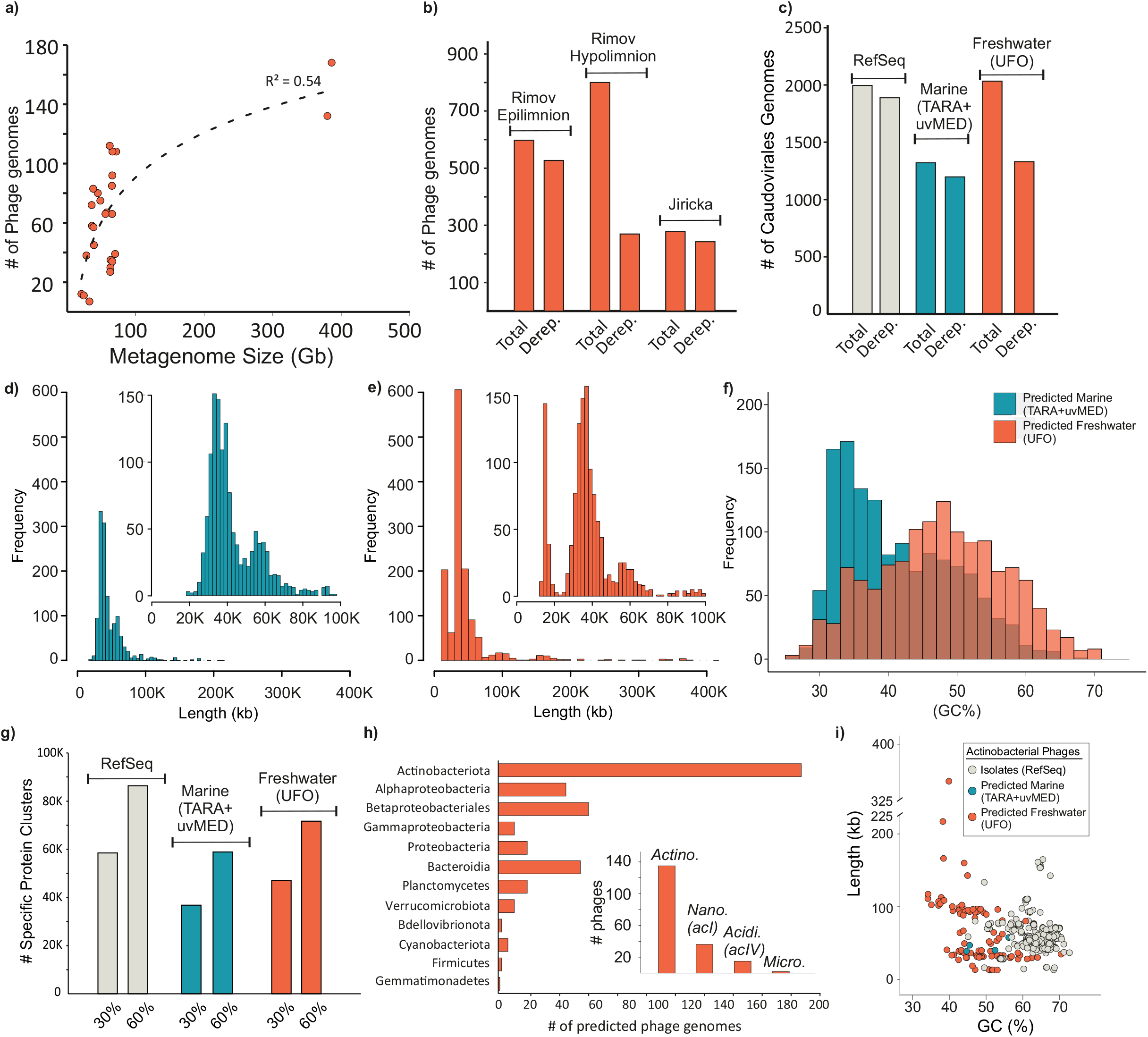
Genome statistics. a) Complete phage genome recovery as a function of metagenome size. b) Total number of phage genomes vs dereplicated genomes from Řimov reservoir (epi- and hypolimnion) and Jiřicka Pond. c) Comparison of all phage genomes and dereplicated phage genomes from Viral Refseq, Marine habitat (TARA+uvMED) and the freshwater habitat (UFO). d) Length distribution of marine phage genomes (inset: enlarged view of phages up to length 100K). e) Length distribution of recovered freshwater phage genomes (inset: enlarged view of phages up to length 100K). f) Comparative GC% distributions of marine and freshwater phage genomes g) Number of unique protein clusters in RefSeq, Marine and Freshwater phage genomes at 30% and 60% identity. h) Number of phage genomes with predicted host of the freshwater dataset (inset: the same for lower taxonomic ranks within Actinobacteria; Actino: Actinobacteriota; Nano: Nanopelagicales; Acidi: Acidimicrobiales; Micro: Microbacteriales) i) GC% vs length for predicted (freshwater, marine) and known actinobacterial phages (RefSeq).

### Phage genome analyses

A total of 598 complete phage genomes were recovered from the Římov epilimnion (10 samples), 800 from the hypolimnion (8 samples) and 279 from Jiřická (5 samples). Upon dereplication (genomes with >95% identity and >95% coverage treated as one, see methods), these numbers reduced by nearly three-fold for the hypolimnion suggesting repeated capture of nearly identical genomes from multiple samplings. We found only a single instance of a phage that was nearly identical in two habitats (Římov and Jiřická).

The comparison of recovered freshwater phage genomes to representative sets of phages from Viral RefSeq (1996 genomes) and the marine habitat (1335 genomes) [23–25] is shown in Fig. 1c. Intriguingly, the genome size distributions of marine and freshwater phage genome sizes appear similar, except for a pronounced peak at small genome size (ca. 15 Kb) in the freshwater datasets. We recovered 155 “miniphage” genomes that were <15 Kb in length (minimum length 13.5 Kb). This somewhat bimodal distribution is remarkably reminiscent of cell size distributions of prokaryotes themselves in freshwater [26] and while not conclusive in itself, this suggests that phage size distribution mirrors host cell size. That such peaks are not visible in the size distributions from isolate phages (Supplementary Figure S3) also points towards a more ecological explanation for the bimodal distribution in freshwater datasets. On the other hand, we also recovered 27 “megaphage” genomes (>200 Kb in length, maximum length 446 Kb) that are similar in genome size to some recently described phages from the human gut microbiome [27].

Common to both freshwater and marine habitats, two frequently recovered genome sizes appear to be ca. 40 Kb and 60 Kb, with 40 Kb being the most frequent. Unsurprisingly perhaps, genomic GC% of recovered phage genomes mirrors the GC% of the habitats (Figure 1f and Supplementary Figure S4). As a measure of how many novel proteins are available in our freshwater phage dataset, we clustered all proteins in these datasets at two percentage identity levels (30% and 60%) [28]. The RefSeq dataset has the maximum number of unique protein clusters (not found in the others), followed closely by the UFO dataset (Figure 1g). Additionally, the number of distinct Pfam domains detected in each dataset were 1761, 927 and 932 for RefSeq, marine and freshwater datasets respectively. These statistics suggest the UFO complete phage genome dataset adds significant novelty in phage sequence space. We also applied vContact2 (that uses Viral RefSeq phage genomes as references) to assess the novelty of the recovered phages [29]. Of the 1330 freshwater and 1202 marine phage genomes (both dereplicated), 775 freshwater phages could not be assigned to any known RefSeq or marine phage cluster, thus remaining either unclassified or clustering only with freshwater phages. Nearly half of marine phages (n=553) also did not cluster with any freshwater or RefSeq phage suggesting the existence of extremely divergent phage populations in these habitats. This is also seen in an all-vs-all comparison of all phage genomes, where the freshwater phages form large clusters that are only weakly related to other known groups (Supplementary Figure S5).

### Host predictions and lifestyle strategies

Using multiple methods (host genes, similarity of tRNA integration sites and presence of CRISPR spacers) we were able to predict hosts for 404 phage genomes (ca. 20%). The maximum number were predicted to be actinophages, largely owing to the presence of the characteristic *whiB* gene (sometimes even in multiple copies, similar to their hosts) that are taxonomically restricted to members of the actinobacterial phylum [19]. These actinophage genomes show an extremely broad size distribution, with multiple “miniphages” (n=15, 6 dereplicated clusters) and “megaphages” (n=3, 2 dereplicated clusters) (Figure 1h). It appears that the most abundant microbes in the freshwater water-column are infected by the full-size range of tailed phages ranging from as small as 14 Kb to as large as 347 Kb.

While for most actinophages we could not specifically pinpoint the host, it was possible for a few, and we predict at least 36 to infect the most abundant lineage in freshwaters (‘*Ca*. Nanopelagicales’, acI lineage [30]). In addition, others that possibly infect freshwater *Acidiimicrobia* (acIV lineage [31]) and *Microbacteraceae* (Luna cluster [32]) were also found (Supplementary Table S2). In comparison to known actinobacterial phages (from cultured isolates), those from the metagenomes display a wide range of GC content and lengths (Figure 1f), with a somewhat lower genomic GC% corresponding to the relatively lower GC% of freshwater *Actinobacteria* (esp. ‘*Ca*. Nanopelagicales’, 42% GC, Neuenschwander et al. 2018).

We also present the first phages predicted to infect multiple other abundant freshwater groups e.g. *Betaproteobacterales, Alpha-* and *Gammaproteobacteria*, *Bacteroidetes*, *Planctomycetes*, *Cyanobacteria*, *Gemmatimonadetes*, etc. (Figure 1h, Supplementary Figure S5, Supplementary Table S2). We recovered only six recognizable cyanophages from our freshwater datasets. We attribute this to the high abundances of filamentous *Cyanobacteria* in freshwaters that are excluded by our filtration and phage recovery methodology. Amongst the other predicted hosts are several well-known freshwater genera (e.g., *Limnohabitans*, *Polynucleobacter, Fonsibacter, Flavobacterium, Novosphingobium, Sphingomonas*, etc. Supplementary Table S2) with no described phages so far, except for ‘*Ca*. Methylopumilus’ [13].

Remarkably, 24 freshwater phages were found to encode the toxin ADP-ribosyltransferase. These are eukaryotic toxins related to VIP2 like toxins (a special class of AB toxins), that inhibit actin polymerization [33] and have been previously suggested to function as a “trojan horse”, effecting targeted killing of eukaryotic predators that phagocytose phage-infected microbes [19,34,35] (Figure 2). These are frequently encoded in phage genomes, and have been found in both free-living phages or inserted prophages, e.g. Shiga toxin [36]. We also found evidence of the same toxin in six marine phage genomes and 116 phages from RefSeq infecting isolates of *Escherichia*, *Mycobacterium*, *Aeromonas*, etc. (Supplementary Table S3). Some of the freshwater phages encoding this toxin are predicted to infect Actinobacteria. Recently, phages have been shown to also encode ribosomal proteins that likely assist phage protein translation during infection (Mizuno et al 2019). We found 20 freshwater phages encoding at least one ribosomal protein (either S21 or L12) (Supplementary Table S4).

**Figure 2.**
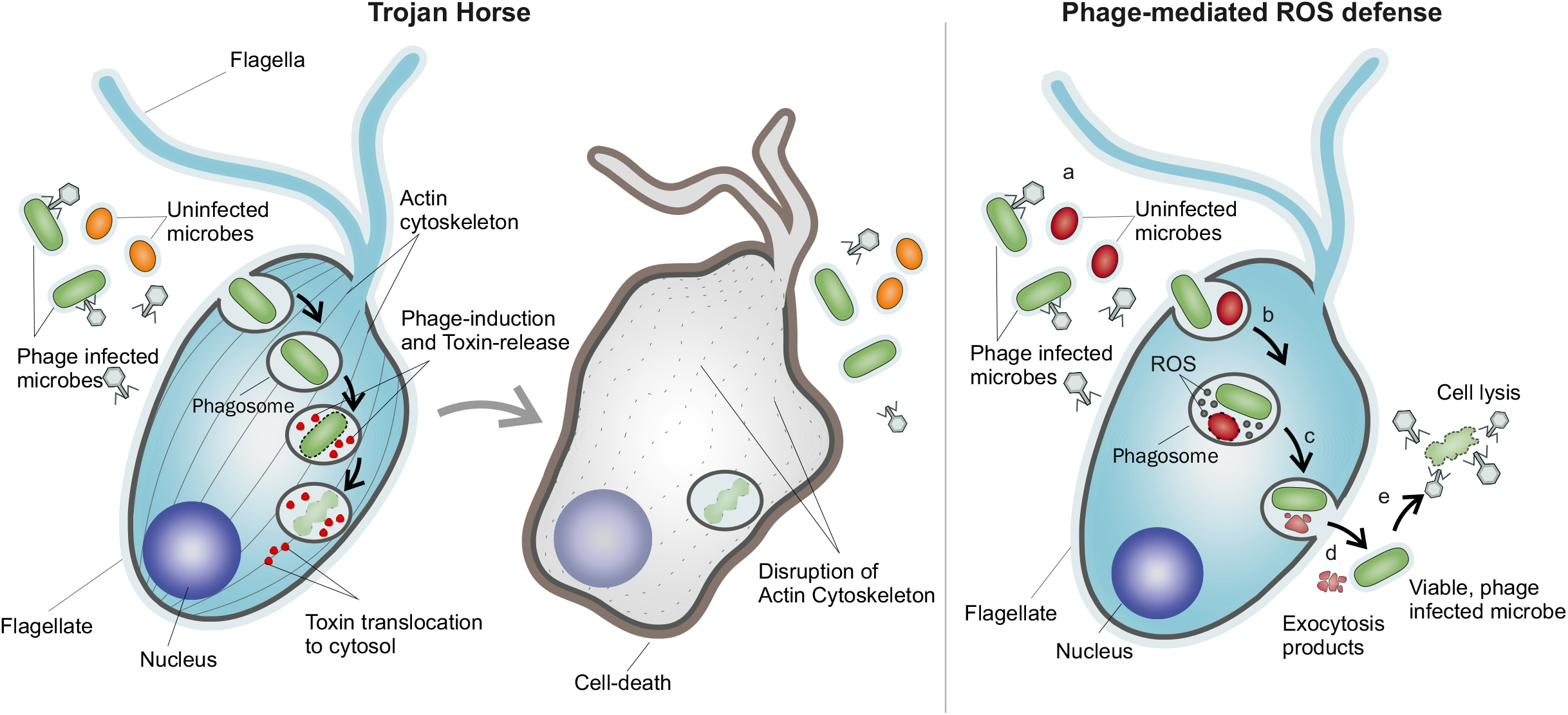
Viral life strategies in freshwater environments. Left: The trojan horse strategy. A phage encoding the eukaryotic toxin ADP-ribosyltransferase (VIP2 family) infects a microbe that is then ingested by a eukaryotic predator (a flagellate is shown). In the phagolysosome the toxin is expressed and released after cell lysis into the phagolysosome from where it translocates to the cystosol. In the cytosol the toxin inhibits actino polymerization leading to cell death. Right: Phage-mediated ROS defense. (a-b) A phage encoding ROS-defense genes e.g. thioredoxin, glutaredoxin is ingested by a flagellate. (c-d) The damage produced to the microbe by the ROS present in the phagolysosome will be reduced by the expression of phage encoded genes, which will favour survival during the oxidative burst. (e) When the phage infected microbe is released outside, cell lysis by the phage can proceed.

More than 10% of phages (i.e. 254 phage genomes) harbored genes involved in oxidative stress mitigation (Supplementary Table S5). Aquatic bacteria routinely experience the damaging effects of reactive oxygen species (ROS) (superoxide, hydrogen peroxide and hydroxyl radicals) produced by their own metabolic machineries, released by other community members or generated by UV-induced photochemical reactions [37]. In defense, bacteria employ several strategies to prevent ROS formation, deactivate ROS and repair the ROS-induced damage. The widespread occurrence of ROS defense mechanisms in bacteria designates oxidative stress as one of the main threats to their fitness and a major culprit in mortality [37–39]. Thus, based on the plethora of ROS-defense mechanisms found in these freshwater phages e.g. ferritin prevents ROS formation; superoxide dismutases and glutathione peroxidases inhibit ROS; thioredoxins and glutaredoxins repair oxidized amino acids particularly cysteine and methionine and PAPS reductases boost reduced sulfur group assimilation [39,40] (Supplementary Table S5), we consider that phages could provide their hosts the means to combat the harmful effects of oxidative stress. Such a strategy could be beneficial for phages, as it ensures the survival (during the lytic cycle) and proliferation of the hosts (during the lysogenic cycle) and protects their own proteins and DNA against oxidative damage. Given that nearly 10% of all recovered phages encode some ROS defense genes, and the high rate of infections in the natural environments (estimated to be up to ca. 25% [41]), it also appears that this strategy is commonly employed. However, unicellular eukaryotes (primarily flagellates) may consume up to 50% of bacteria in freshwater habitats on a daily basis [42]. This significant number, coupled with high phage infection frequencies suggests that multiple phage-infected bacteria must be ingested by these flagellates. Not all microbes are fully digested in the food vacuoles and many are expelled outside again [43,44]. It is quite likely that ROS-defense mechanisms improve the odds for surviving the phagolysosome where reactive oxygen species are discharged to destroy bacteria (oxidative burst). We postulate that a bacterium infected by a phage containing a ROS-defense mechanism will have a selective advantage during phagocytic flagellate grazing. Thus, the phage-encoded proteins could help the host survive the high ROS environment that characterizes the phagolysosomes [45], i.e. a phage -mediated ROS defense (Figure 2). In both these mechanisms, (the trojan horse and phage-mediated ROS defense), genes that enhance the preservation of the host-phage interaction at the expense of the eukaryotic predators appear to be selected and widely encoded in environmental phage genomes.

### Phage abundance time-series

Owing to the samples from multiple time points in the epi- and hypolimnion of Římov reservoir, we were able to recover several, nearly identical phage genomes repeatedly. The most extreme case was that of a predicted actinophage that was recovered twelve times, in distinct seasonal phases (spring bloom, summer and winter). Remarkably, even though it was retrieved at different times of the year, only seven “variant” locations are seen (Supplementary Figure S6). Six of these are present in hypothetical genes and one in an intergenic region. Exhaustive sequence searches using jackhmmer [46] and HHpred server [47] revealed little clues to their functions. However, the retrieval of multiple, nearly identical phage genomes from multiple time-points and strata suggests that some lineages are persistent, likely also owing to the constant presence of the host, in this case, *Actinobacteria* that are always abundant in the Římov reservoir [20] (Supplementary Figure S2). Similar to this phage, we also recovered multiple other examples that remain unchanged during our sampling efforts (Supplementary Table S2).

Remarkably, the abundance patterns of phages recovered from the Římov reservoir (n=1398) (Figure 3) suggest seasonality in their appearance. Distinct sets of phages peak in different layers during summer stratification (epi- or hypolimnion) or mixis (both spring and early winter). In the hypolimnion, some phages are persistent throughout the year while others appear only during stratification. Moreover, the abundances (coverage per GB) for each phage across the entire timeline of the Římov reservoir (18 samples) show that most phages in the epilimnion transiently achieve high abundances followed by near disappearance (a boom and bust scenario) while several in the hypolimnion appear to be more persistent and are recovered at multiple time points (Figure 3). The shorter timeline of the Jiřická samples also shows a near continuous replacement of the abundant phages comparable to the Římov epilimnion (Supplementary Figure S7) in the relatively warm time period sampled here. However, as this time series is far shorter (lasting only a few summer months, Supplementary Figure S2), it remains to be seen if in such dynamic systems anything resembling persistence as observed in the colder hypolimnion of the Římov reservoir.

**Figure 3.**
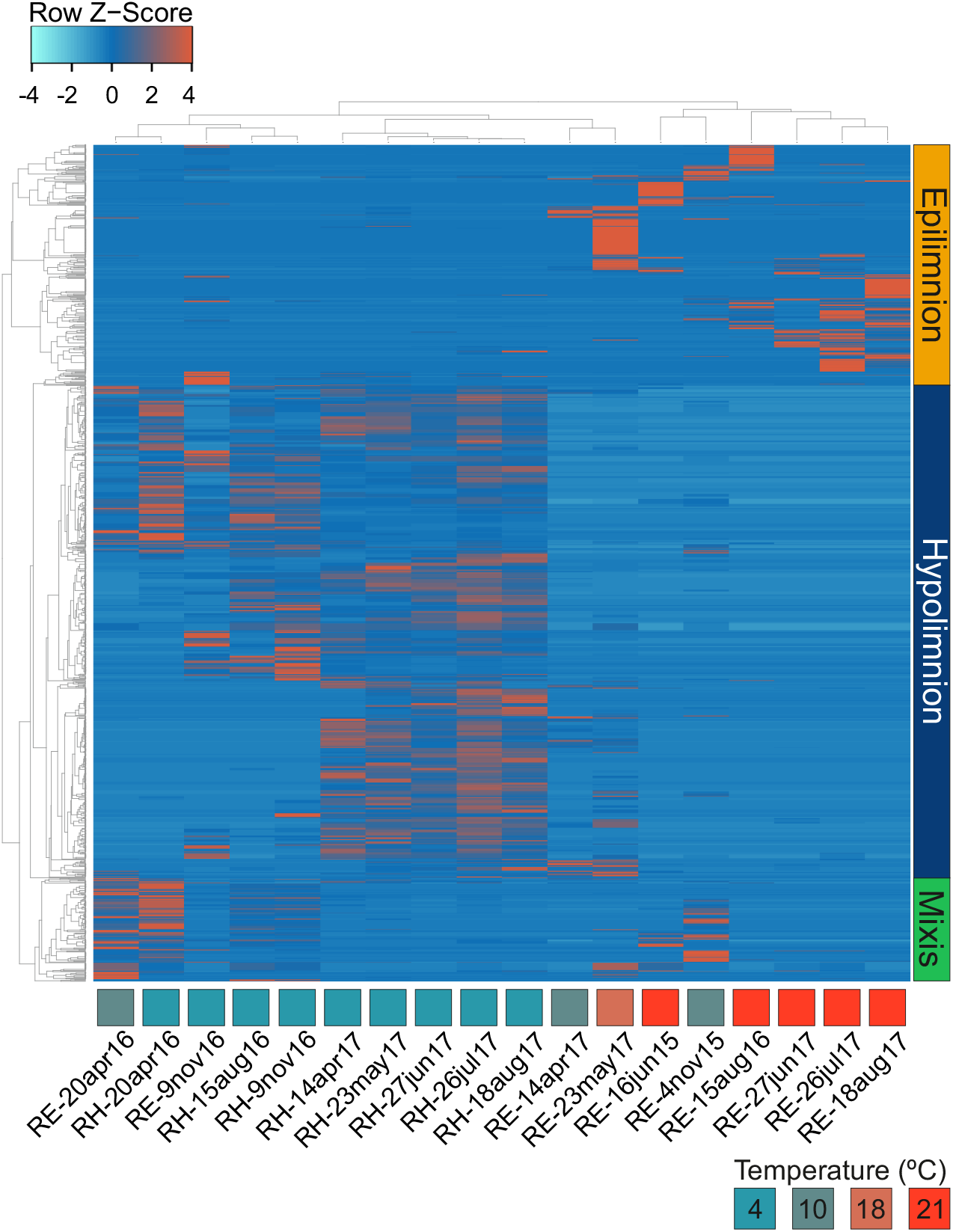
Relative abundance of 1398 Řimov phages in 18 metagenomes. A heatmap of abundances is shown (coverage/Gb of metagenome normalized by Z-score). Phages are clustered by sample and abundance (average linkage, Spearman Rank correlation). Vertical bars on the right side indicate the classification of the clusters. Columns are annotated with the temperature, depth of the sample and the sampling date (RE = 0.5m and RH = 30m). Temperature color key is shown at top right.

### Spatiotemporal dynamics of actinophages and their hosts

As actinobacterial phages were the largest identifiable group (owing to the presence of the *whiB* gene), and that *Actinobacteria* are known to be dominant members of the community throughout the year (Supplementary Figure S2), we chose to focus subsequent analyses on both recovered actinophages and actinobacterial MAGs. Abundance profiles of all recovered actinophages encoding *whiB* (confident predictions, n=125) are shown in Figure 4. Phages that are nearly identical, i.e. persistent phages (>95% identity and >95% coverage) are shown as part of a cluster. The profiles within a cluster appear homogenous but different clusters show distinct preferences for either the epilimnion or the hypolimnion, suggesting that their hosts (*Actinobacteria*) would also show similarly distinct patterns.

**Figure 4.**
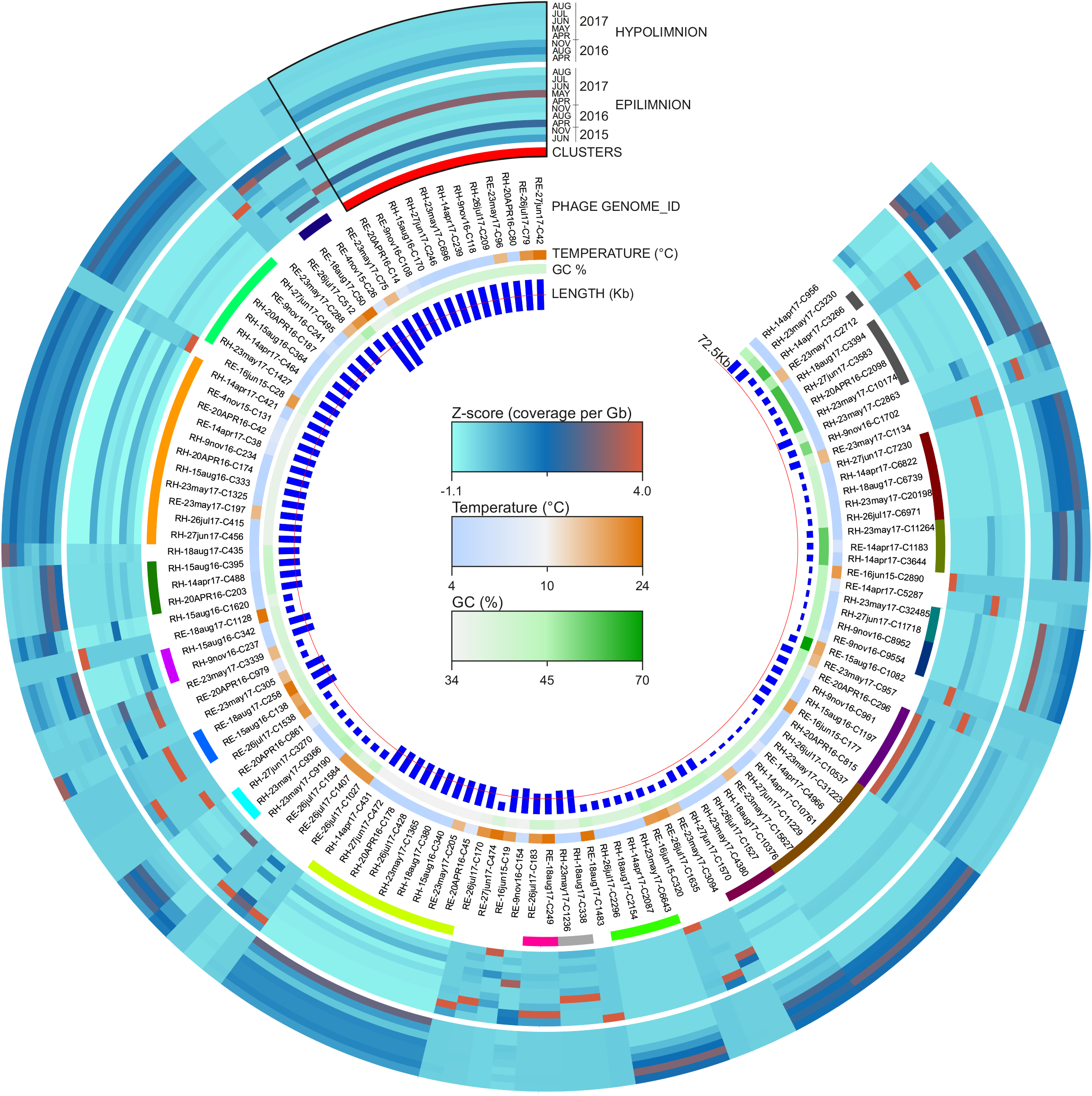
Abundance profiles of the WhiB encoding actinophages recovered from Řimov reservoir. Actinophages (n=125) are arranged based on average amino acid identity (dendrogram not shown). From inside out the rings represent: phage genome size in Kb (red line indicates the average genome length, 72.5Kb), phage genome GC%, water temperature of samples whereof phages were assembled, span of clusters of phage genomes grouped by similarity (nucleotide identity >95%), abundance profiles of each actinophage in the epi-and hypolimnion time series datasets of Řimov reservoir (coverage per Gb of metagenome normalized by Z-score). Color keys are shown in the center. Phages of Cluster1 are outlined in black (top left of the circle).

We recovered 444 actinobacterial metagenome-assembled genomes (MAGs) from the Římov Reservoir datasets, whereof 305 MAGs fulfilled the criteria for further analyses (see Methods). A phylogenomic analysis of the recovered actinobacterial MAGs in context of known isolate genomes is shown in Figure 5. Most MAGs are placed within the three known groups of *Actinobacteria* frequently found in freshwater habitats, ‘*Ca*. Nanopelagicales’ (n=280), *Acidimicrobiia* (n=114) and *Microbacteriaceae* (n=37) (Supplementary Table S6). In particular, the order ‘*Ca*. Nanopelagicales’ are the most cosmopolitan microbes from freshwater [30,48,49] and have only recently been brought into culture [30,50]. Isolates from *Microbacteriaceae* are also available [32,51] but no cultured representatives exist yet for *Acidimicrobiia* and these are described only from metagenome-assembled genomes [52]. Within the order ‘*Ca*. Nanopelagicales’, two genera are defined, ‘*Ca.* Planktophila’ (formerly acI-A) and ‘*Ca.* Nanopelagicus’ (formerly acI-B) [30,53]. However, several other lineages have been described from 16S rRNA based surveys, e.g. acI-C [53], acSTL [54], acTH1 [55]. While recently a single genome from acI-C isolate has become available [50], no isolates or genomes have been described for either acSTL or acTH1. Based on the presence of 16S rRNA sequences, we identified MAGs that belong to acI-C2, acSTL, and acTH1 lineages. The acI-C2 related MAGs branch outside the acI-A and acI-B in accordance with known phylogeny. Both acSTL and acTH1 appear as a deep-branching sister group to other ‘*Ca*. Nanopelagicales’ but still belong to the same order (as classified by GTDB [56]). In addition, we also recovered ‘*Ca*. Nanopelagicales’ MAGs that are basal to all known groups (Figure 5).

**Figure 5.**
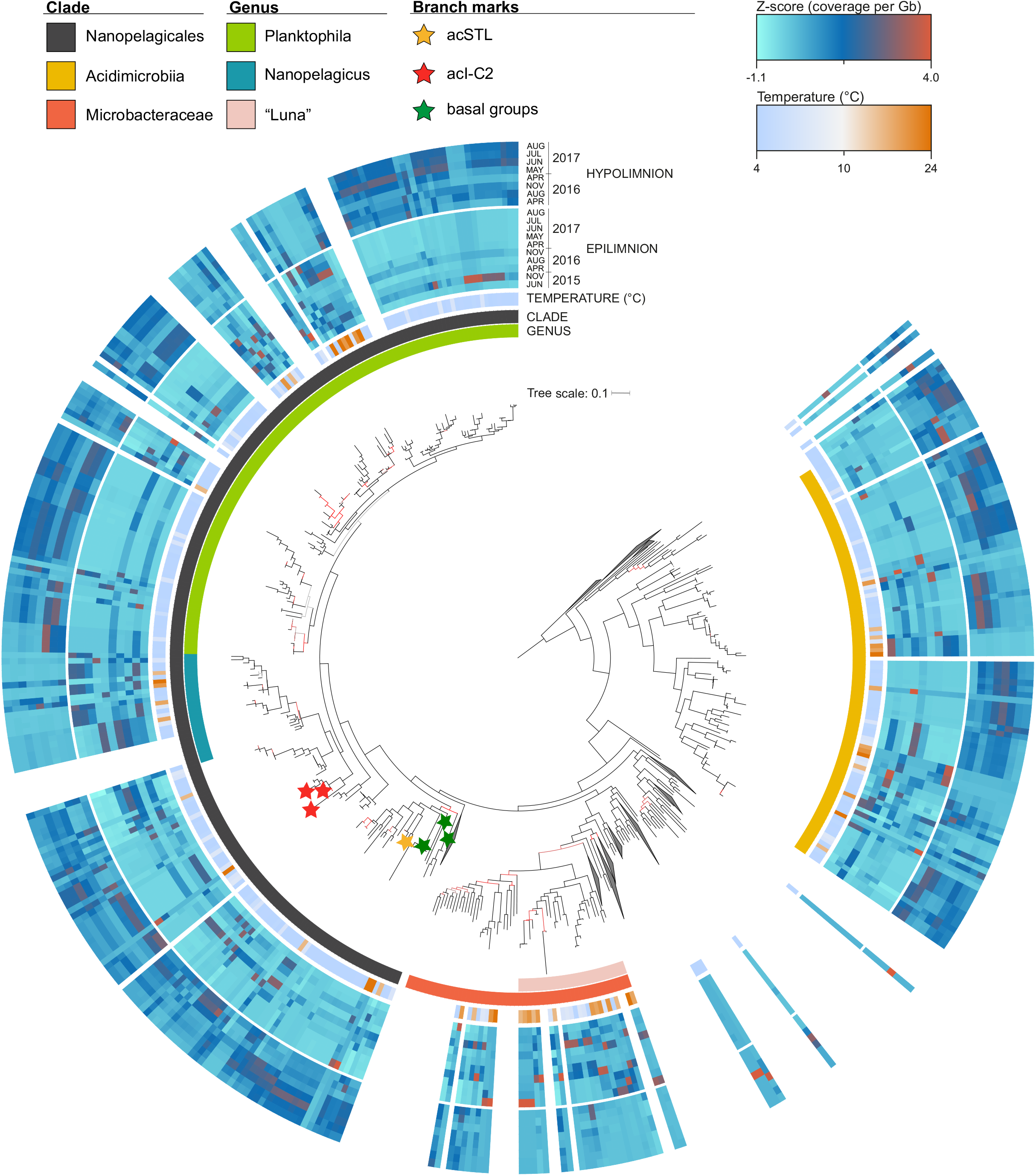
Abundance profiles of actinobacterial genomes and MAGs combined with phylogenomic analysis. The tree was obtained using complete reference genomes (n=205) and recovered MAGs (n=350) of Řimov reservoir time series metagenomes (see methods). From inside out the rings represent: the larger ring covering some tree branches highlights detailed taxonomic levels of Microbacteraceae and Nanopelagicales, the second ring shows class or order of Actinobacteria, followed by water temperature of samples whereof MAGs were assembled and abundance profiles for each MAG in the epi- and hypolimnion time series datasets of Řimov reservoir (coverage per Gb of metagenome normalized by Z-score). The abundance profiles are shown only for MAGs recovered from the Řimov reservoir. The red and yellow stars indicate acSTL and acTH1 MAGs with 16S rRNA sequences respectively. Green stars indicate deep branching basal groups within the order Nanopelagicales. Branch colors reflect bootstrap support (UFboot): black: >=95, red: 45-95 and gray: <45. Color keys are shown at the top.

The abundance profiles of these *Actinobacteria* MAGs revealed that the vast majority are more abundant in the hypolimnion than the epilimnion, especially ‘*Ca*. Nanopelagicales’ and *Acidimicrobiia*, while the reverse is true for *Microbacteriaceae*, that are nearly always more abundant in the epilimnion (Figure 5). MAGs that are in close phylogenetic proximity (for instance within the genus ‘*Ca*. Nanopelagicus’) do not necessarily show similar abundance patterns implying niche divergence even within closely related organisms (Neuenschwander et al. 2018) and several show only peaks in epilimnion or hypolimnion alone. Remarkably, the temporal abundances of both phages and their hosts mirror three distinct states of the reservoir, warm and stratified epilimnion, cold hypolimnion and mixed water column. This is seen even more clearly in the predicted actinophages (Figure. 4) and reflected in the abundances of their hosts (Figure 5).

The sporadic peaks of phages observed in the epilimnion reflect the transient niche that it truly is, in comparison to the apparently more stable environment of the hypolimnion (with little temperature variation). The ground state for Římov is a low-temperature regime, lasting for nearly two-thirds of the year. Only at the end of the spring overturn until onset of winter a shallow, peripheral zone with higher temperature and light intensity (i.e. epilimnion) is established within which periodic blooms of photosynthetic organisms are observable [57]. Regardless of the sporadic peaks, we captured a complete genome of an actinophage in summer (at high abundance), and the same genome was recovered repeatedly at lower abundances, from both epi- and hypolimnion samples and at widely different temperatures (4ºC to 24ºC) at different times of the year (cluster 1 in Figure 4). This suggests its host experiences conditions favorable for its increased abundance in the warmer epilimnion what leads to the higher abundance of its phage. The persistent recovery of such a phage also suggests that its host remains available throughout the year. In line with these observations, many actinobacterial MAGs show short-lived maxima in the epilimnion followed by lower abundances in the hypolimnion as observed earlier via fluorescence in situ hybridization with species to genus-specific probes [30]. It has also been shown recently that in the absence of the optimal host phages may switch to sub-optimal hosts [58], further driving diversification and likely helping extend the longevity of phage lineages.

The major observable dynamic in the hypolimnion is the remarkably similar abundance patterns of the hosts and their phages, i.e. both show persistence from the onset of stratification till next spring. It appears that the hypolimnion maintains a large pool of highly related host genomes most of which are well-adapted to the long-lasting low-temperature regime of the reservoir. A fraction of these might find favorable niches in warmer temperatures, blooming, and then retreating within the hypolimnion at winter onset. However, even with the observable “persistent” abundances of phages, the hypolimnion is not without its perturbations (not in temperature but in other environmental variables, e.g. hypoxia, irregular nutrient input). Less obvious, but sporadic peaks are observed in the hypolimnion as well (Figure 3), what would suggest clonal expansions of the host, as has been reported for some *Planctomycetes* [59], suggesting similar dynamics are also played out in deeper waters.

## Conclusions

Freshwater habitats are relatively accessible to monitoring and the use of time-series metagenomes allows sensitive, genome-based surveys to capture both recurrent and anomalous changes in community composition. In this work, we used a deep-sequencing time-series approach to viral ecology aimed towards the recovery of a representative freshwater phage genome collection that significantly expands the known viral sequence space and revealed viruses infecting many different freshwater phyla for which none were known before. This collection of complete phage genomes should serve as a critical reference in boosting environmental genomics of viruses in freshwater habitats at large, illuminating their wide genomic repertoire and revealing their ecological interactions not only with their microbial hosts, but also with the predators of their hosts, i.e. unicellular eukaryotes.

While this study was focused only upon phages, the metagenomes described here are expected to shed light on many other viral groups as well e.g. ssDNA viruses, phycodnaviruses and virophages. With the availability of relatively inexpensive and increasingly higher throughput in sequencing technologies and the advent of even longer reads, freshwater viral ecology can transition from a gene-based to a genome-centric view of the viral world around us.

## Data Availability

Sequence data for all metagenomes generated in this work are archived at DDBJ/EMBL/GenBank and can be accessed under the Bioproject PRJNA429141 for the Římov reservoir and Bioproject PRJNA429145 for Jiřická pond. All phage genomes and the actinobacterial metagenome assembled genomes are available in NCBI Bioproject PRJNA449258.

## Acknowledgments

The authors thank Petr Znachor and Pavel Rychtecký for help with sampling of Římov Reservoir and Petr Porcal for sampling Jiřická Pond. VSK was supported by the research grant 17-04828S (Grant Agency of the Czech Republic). A-Ş. A was supported by the research grants 17-04828S (Grant Agency of the Czech Republic) and MSM200961801 (Academy of Sciences of the Czech Republic). MM was supported by the Postdoctoral program PPPLZ (application number L200961651) provided by the Academy of Sciences of the Czech Republic. MMS was supported by research grant 19-23469S (Grant Agency of the Czech Republic). RG was supported by the research grants 17-04828S (Grant Agency of the Czech Republic) and CZ.02.1.01/0.0/0.0/16_025/0007417 (ERDF/ESF).

## Methods

### Sampling site and collection

*Site I:* Římov reservoir (Czech Republic, 48.846361 N 14.487639 E) with 2.06 km^2^, volume 34.5×10^6^ m^3^, length 13.5 km, circum-neutral pH, maximum depth 40m, average depth 16.5m, retention time ~100 days, dimictic, meso-eutrophic, moderately humic). Built in 1979, on the Malše River, it is part of the Czech Long Term Ecological Research network [22,60,61]. Eighteen water samples were collected from epilimnion (0.5m, n=10) and hypolimnion (30m, n=8) from June 2015 to August 2017. Two of these have been published previously [59,62] and the rest were generated in this study. Vertical profiles of the physicochemical characteristics of the water column (temperature, pH, oxygen; GRYF XBQ4, Havlíčkův Broc, CZ) and chlorophyll *a* (FluoroProbe TS-16-12, bbe Moldaenke, Kiel, Germany) were also taken. *Site II:* Jiřická pond (Czech Republic, 48.616034 N 14.676594 E) with 0.0356 km^2^, volume 6.59 x10^3^ m^3^, pH 5.6-6.2, maximum depth 3.7 m, retention time ~5-7 days, dystrophic, located in the Novohradské mountains of Southern Bohemia [21]. Five samples were collected from the epilimnion (0.5m) from May 2016 to August 2017.

### Filtration and DNA extraction

All water samples (ca. 10 L each) from Římov reservoir and Jiřická pond were sequentially filtered through 20 µm, 5 µm and 0.22 µm polycarbonate membrane filters (Sterlitech, USA). The 0.22 µm filters (containing the 5 - 0.22 µm microbial size fraction) were cut in small pieces (≅ 3–5mm) using sterile scissors and processed for DNA extraction using the ZR Soil Microbe DNA MiniPrep kit (Zymo Research, Irvine, CA, USA), following the manufacturer’s instructions.

### Preprocessing of metagenomic datasets

Shotgun sequencing was performed using Illumina HiSeq4000 (for samples of 2015 and 2016 −2x 151bp) (BGI HongKong, China) and Novaseq 6000 (for samples of 2017 and Jiřická 2016 - 2x 151bp) (Novogene, HongKong, China). Raw Illumina metagenomic reads were preprocessed in order to remove low-quality bases/reads and adaptor sequences using the bbmap package [63]. Briefly, the PE reads were interleaved by *reformat.sh* and quality trimmed by *bbduk.sh* (using a Phred quality score of 18). Subsequently, *bbduk.sh* was used for adapter trimming and identification/removal of possible PhiX and p-Fosil2 contamination. Additional checks (i.e. *de novo* adapter identification with *bbmerge.sh*) were performed in order to ensure that the datasets meet the quality threshold necessary for assembly. The preprocessed reads were assembled independently with MEGAHIT (v1.1.5) [64] using the k-mer sizes: 49,69,89,109,129,149, and default settings.

Publicly available freshwater metagenomes (total 149 datasets) were downloaded and assembled as described above. Basic metadata (sampling date, location, depth, Bioproject identifiers, SRA accessions), and sequence statistics of all metagenomes generated or used in this study are provided in Supplementary Table S1.

### 16S rRNA abundance based taxonomic classification

20 million reads were randomly sampled from each metagenome and compared to the SILVA database (version 132) [65] to identify candidate 16S reads using an e-value cutoff of 1e-3 using MMSeqs2 [66]. The candidate reads were further screened with ssu-align [67] to find bona-fide 16S rRNA sequences. The 16S rRNA sequences were compared to the SILVA database using blastn and taxonomy of the best hits was used to obtain the final taxonomic classification.

### Gene prediction, phage detection and annotation

Prodigal was used for gene prediction in metagenomic mode [68]. To retrieve complete phage genomes we selected assembled contigs >10 Kb that provided evidence of a circular genome as described before [19] (n=3576 circles). These sequences were scanned with the VirSorter tool (default settings, Virome and RefSeq decontamination mode, scores 1 and 2) (https://de.iplantcollaborative.org/de/) [69] and MARVEL [70], and those contigs that were detected by either method to be of phage origin were retained. Finally, a set of 2034 genomes were judged to be complete. Existing marine phage genomes were recovered from the Tara Oceans metavirome assemblies[24,25] and the uvMED dataset[71] ans re-run through VirSorter (using the same criteria as for freshwater phage genomes). Additional manual curation was performed using NCBI Batch CDD server to minimize errors in phage identification [72].

### Recovery of actinobacterial genomes

The curated metagenomic datasets of Římov reservoir and Jiřická Pond were mapped using bbwrap.sh [73] (kfilter=31 subfilter=15 maxindel=80) against the assembled contigs (longer than 3 Kb) in a lake-dependent fashion. The resulting BAM files (324 for Římov Reservoir, 25 for Jiřická Pond, respectively) were used to generate contig abundance files with *jgi_summarize_bam_contig_depths* [74] (--percentIdentity 97). The contigs and their abundance files were used for binning with MetaBAT2 [74](default settings). Bin completeness, contamination and strain heterogeneity were estimated using CheckM [75] (with default parameters). Bins with estimated completeness above 40% and contamination below 5% were denominated as metagenome-assembled genomes (MAGs). MAGs were taxonomically classified with GTDB-Tk [56] with default settings. MAGs belonging to Actinobacteria (444), together with reference genomes (205) recovered from public repositories (Supplementary Table 6) were annotated using the TIGRFAMs database [76]. 35 conserved marker proteins (Supplementary Table S7) were extracted from the annotated *Actinobacteria* genomes. MAGs that had more than 19 markers present and reference genomes were used for phylogenetic reconstruction. Briefly, homologous proteins were independently aligned with PRANK [77] (default settings), trimmed with BMGE [78] (-t AA -g 0.5 -b 3 -m BLOSUM30) and concatenated. A maximum-likelihood phylogeny was constructed using IQ-TREE [79] with the VT+F+R10 substitution model (chosen as the best-fitting model by ModelFinder [80]) and 1,000 ultrafast bootstrap replicates [81]. These samples also allowed us to assemble and bin ca. 2400 microbial genomes in order to maximize chances for host prediction for the recovered phages.

### Sequence annotation

All selected viral contigs were compared to the NCBI non-redundant protein database. Protein domains in coding sequences were annotated with Interproscan [82,83]. Searches were performed locally using HMMER3 package [84] with e-value = 1e-3 for Clusters of Orthologous Groups (COGS) [85] and trusted score cutoffs for TIGR Families – TIGRfams [76]. Pfam domains were identified in all datasets using the script pfam_scan.pl (ftp://ftp.sanger.ac.uk/pub/databases/Pfam/Tools/) with the PFAM database release 31 [86].

### Host prediction

Multiple methods were used to assign a host to a phage genome. Host-specific genes were used to predict the host e.g. photosystem genes to link to cyanophages [87] and *whiB* for *Actinobacteria* [19]. Integration sites in the host genome (termed *attB*), usually a tRNA locus, were checked using BLASTN [88] for assigning a specific association between host and phage as described before [71]. CRISPR spacers in microbial genomes were detected using minced (https://github.com/ctSkennerton/minced) and for host prediction spacers were compared to phage genomes using BLASTN with stringent cutoffs (alignment length ≥30 bp, ≥97% nucleotide identity, ≥97% query coverage, ≤1e-5). Additionally a direct comparison of phage genomes to host genomes was made using BLASTN to identify shared nucleotide sequences (alignment length ≥30 bp, ≥97% nucleotide identity, ≥97% query coverage, ≤1e-5) [89].

### Selecting representative phage genomes

*Caudovirales* (tailed phages) phage genomes were divided into three sets, from NCBI Viral RefSeq (n=1996), marine *Caudovirales* (Tara Oceans and uvMED, n=1335) and freshwater *Caudovirales* (n=2034). Within each set all-vs-all blastn comparisons were made retaining significant matches at evalue <1e-3 and >95% nucleotide identity. For each comparison, two phages were considered as belonging to a cluster if the genome coverage of both phages in a pairwise comparison was ≥95%. Clusters of phages that meet these criteria were then merged together if they shared a phage genome in common (single linkage). The longest phage in each cluster was selected as a representative phage genome for this cluster. Clustering at these relatively high nucleotide identity levels and genome coverages is expected to retain phage genomes that are very closely related at the genomic level. At these cutoffs, the number of representatives in each dataset was: RefSeq 1887, marine 1202 and freshwater 1330, making a total of 4419 representative phage genomes.

### Phage proteomic tree

An all-vs-all tblastx comparison was performed for all 4419 phages (-M BLOSUM45 –e 1e-3) and the scores of all significant hits were added together to provide a comparison score for all pairwise comparisons. The comparison scores between two phage genomes were normalized by the self-comparison of both phages to provide a similarity metric (Dice coefficient). For example, 2 * comparison score of A and B / (self-comparison score of A + self-comparison score B). The Dice coefficient was subtracted from 1 to provide a distance measure [25,71]. The resultant distance matrix of the all-vs-all comparison was used to generate 10000 alternative, but equally valid tree topologies using clearcut [90] (clearcut –in distance_matrix.txt –out 10000.trees –n 10000), and a consensus tree was computed using IQ-TREE [79] (iqtree -t 10000.trees –con consensus.tre) that contains confidence values for each comparison. The final tree was visualized in iTOL [91] (http://itol.embl.de).

### vContact2 classification

vContact2 was run on the Cyberverse infrastructure using diamond, freshwater phages only (1330 genomes, dereplicated), marine phages only (1202 genomes, dereplicated), and finally both freshwater and marine phages together (2532 genomes).

### Fragment recruitment

All phage genomes were compared to all metagenomic datasets using RazerS 3 [92] (using cutoffs of >95% identity and alignment lengths ≥50bp) to compute coverage per Gb. For microbial genomes, all rRNA sequences (5S, 16S and 23S) were identified using rrna_hmm [93] and were masked prior to comparisons with metagenomic sequences. At most, 20 million reads were used from metagenomic datasets for computing abundances of microbial genomes while a full set of reads were used for phage genomes. Raw data (coverage per Gb) is given in Supplementary Table S2 (phages) and Supplementary Table S6 (Actinobacteria). A phage genome or microbial genome was considered to be present only when it presented >80% genome coverage. Heatmaps were created using http://heatmapper.ca [94].

## Supplementary Information

**Supplementary Figure S1.**
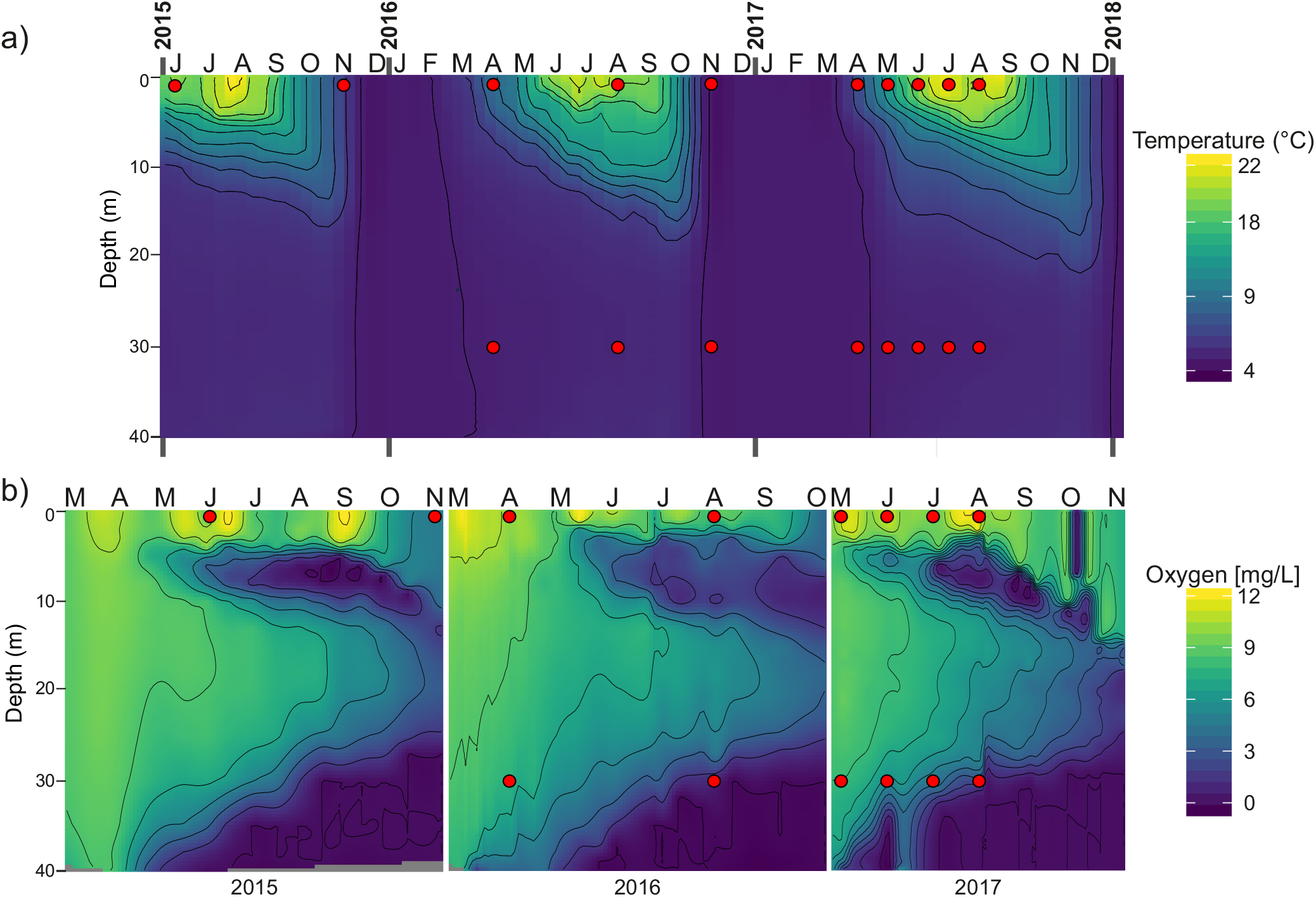
a) Water temperature along a depth profile from June 2015 to December 2017 in Římov reservoir. b) Oxygen concentrations along a depth profile from March 2015 to November 2017 in Římov reservoir. Red circles indicate the time and depth of the samples. Oxygen measurements were not available for all time points.

**Supplementary Figure S2.**
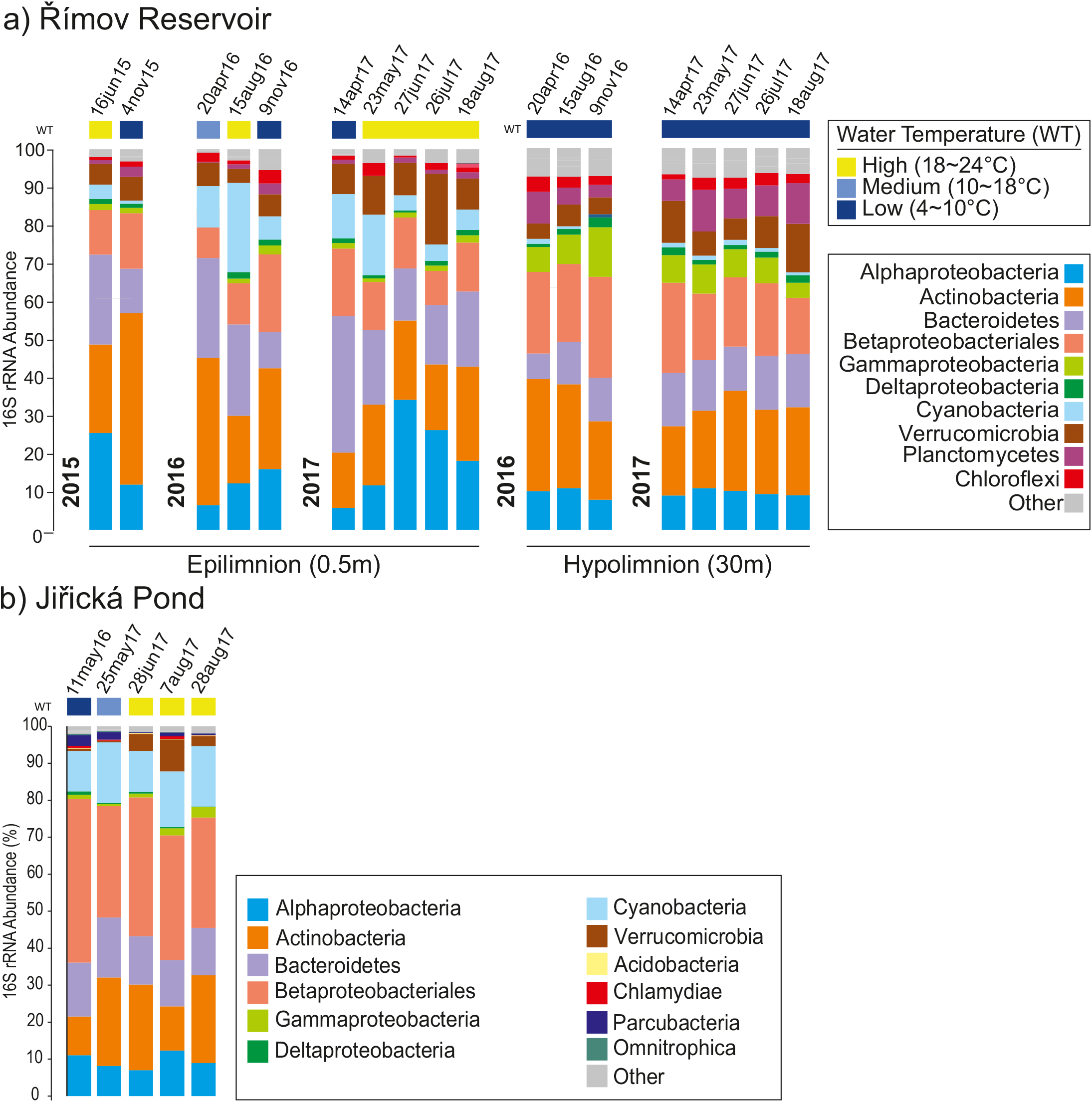
16S rRNA based abundances of prokaryotic groups. The figure depicts the SILVA SSU (Ref NR 99 132) classification of 16S rRNA reads in the metagenomes of (a) Římov reservoir and (b) Jiřická Pond. Water temperature at the time of sampling is also indicated (scale top right).

**Supplementary Figure S3.**
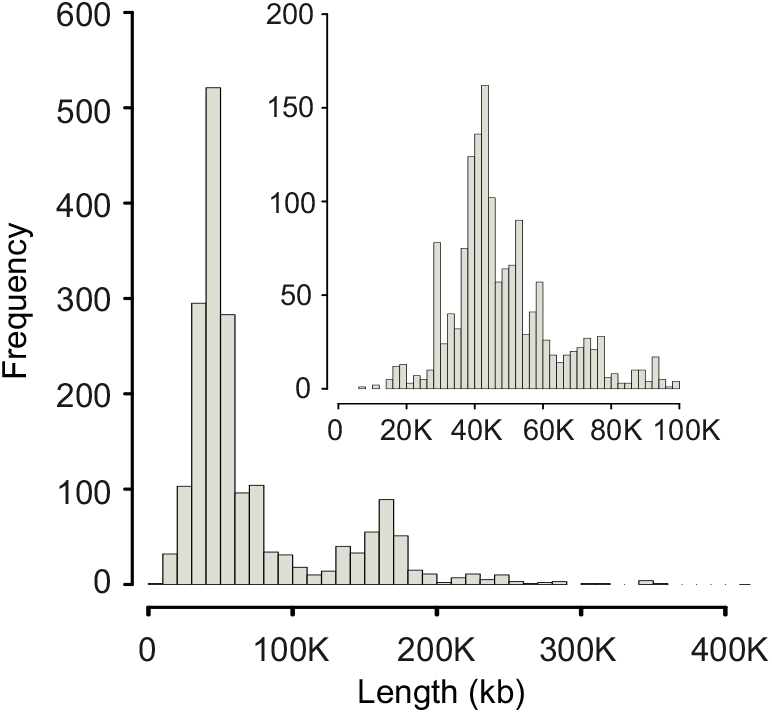
Length distribution of Viral RefSeq phage genomes (inset: enlarged view of phages up to length 100K).

**Supplementary Figure S4.**
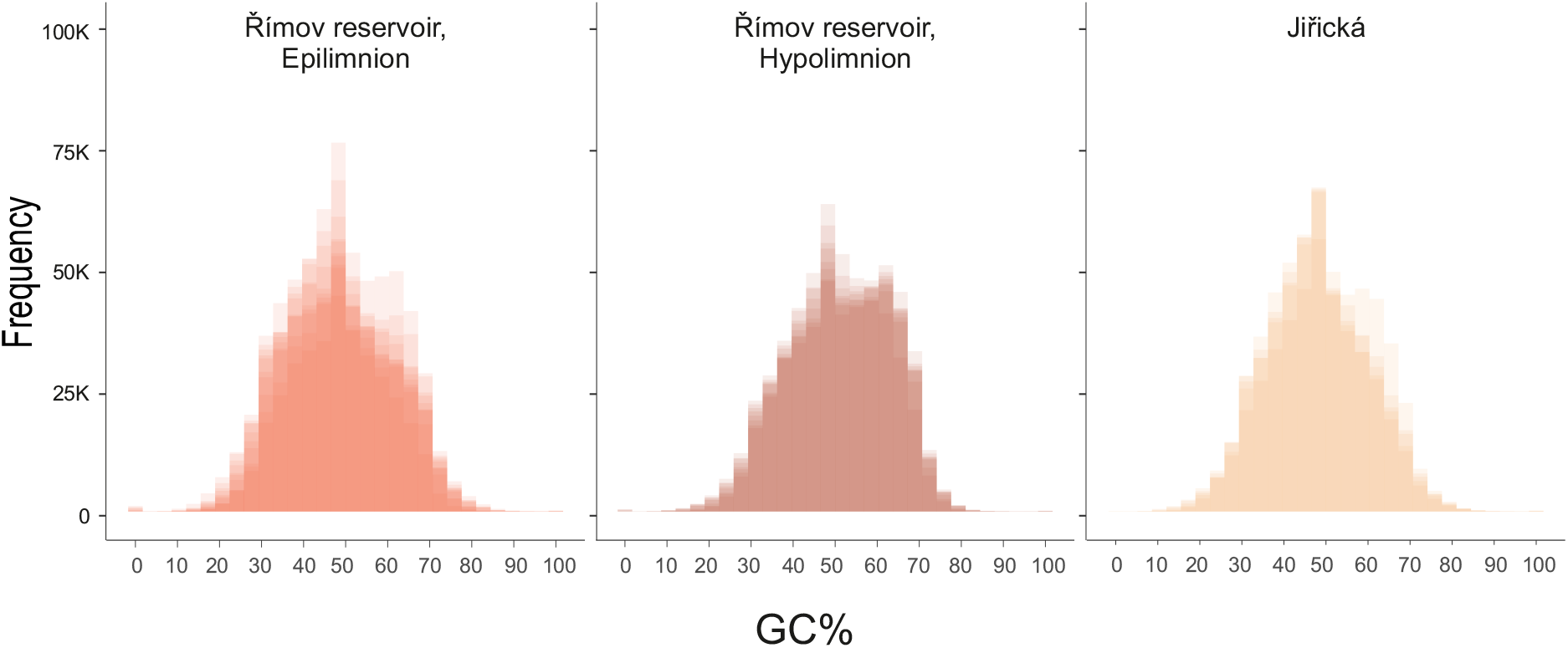
Comparative GC% distributions of freshwater metagenomic data for Římov epilimnion (n=10 datasets), Římov hypolimnion (n=8 datasets) and Jiřická (n=5 datasets).

**Supplementary Figure S5.**
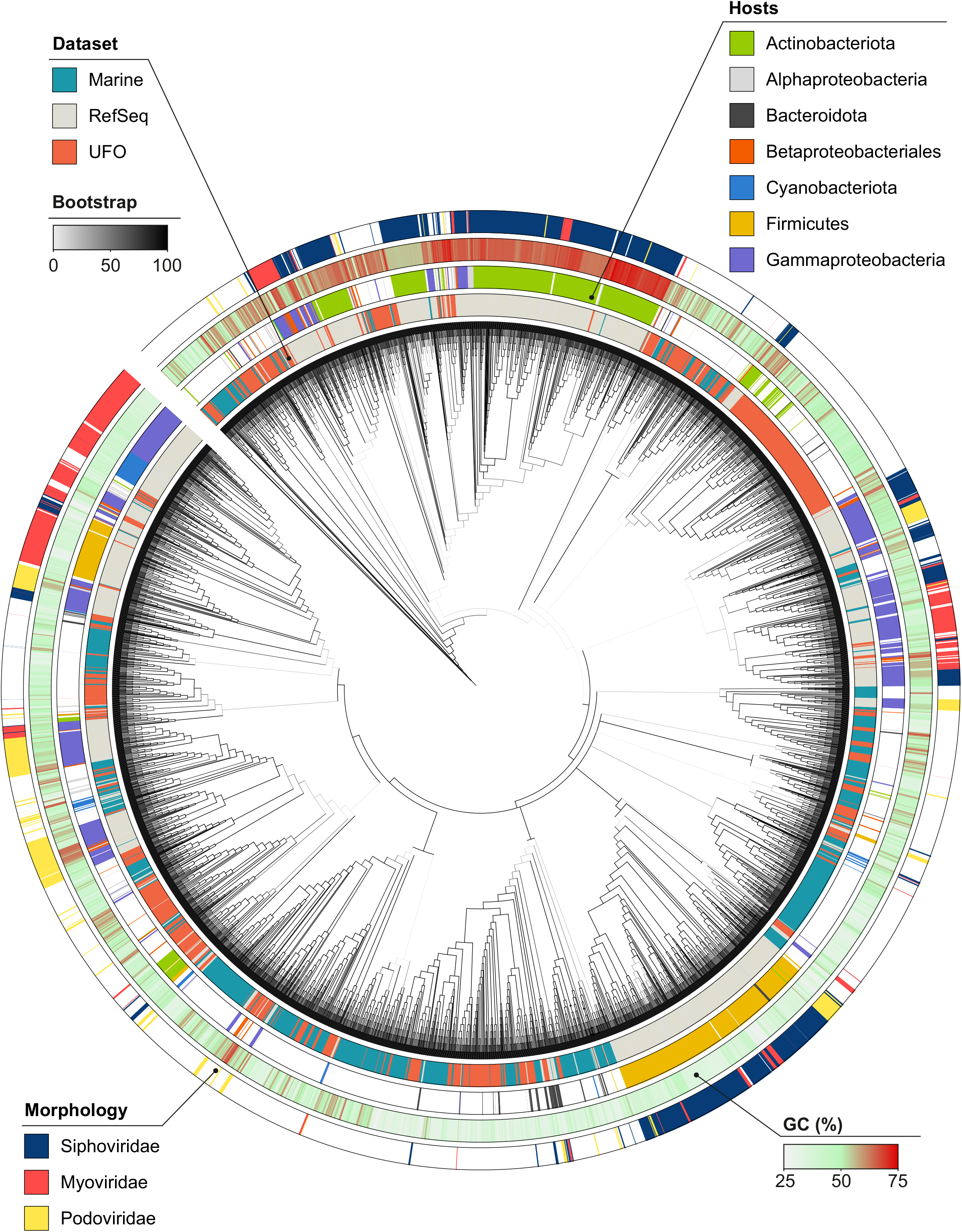
Phage proteomic tree showing relationships of freshwater (n=1330), marine (n=1202) and RefSeq (n=1887) phages used in this study. For more details see methods. From inside out: Gray colored tree branches represent bootstrap values below 100, phage dataset, predicted hosts, genome GC% and known phage morphology (for RefSeq). Figure made with iTOL (http://embl.itol.de).

**Supplementary Figure S6.**
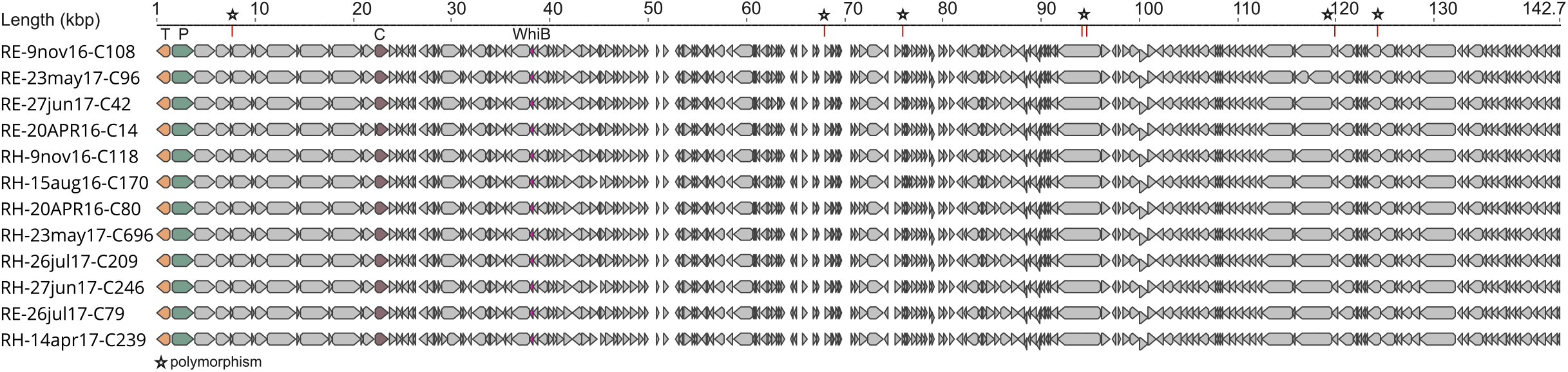
Genomic variations in a persistent actinophage (Cluster1). Whole genome alignment of a phage genome that was recovered independently 12 times. The variant polymorphisms are indicated with a “*” at the top. T: Terminase Large Subunit, C: Major Capsid protein, P: Portal protein.

**Supplementary Figure S7.**
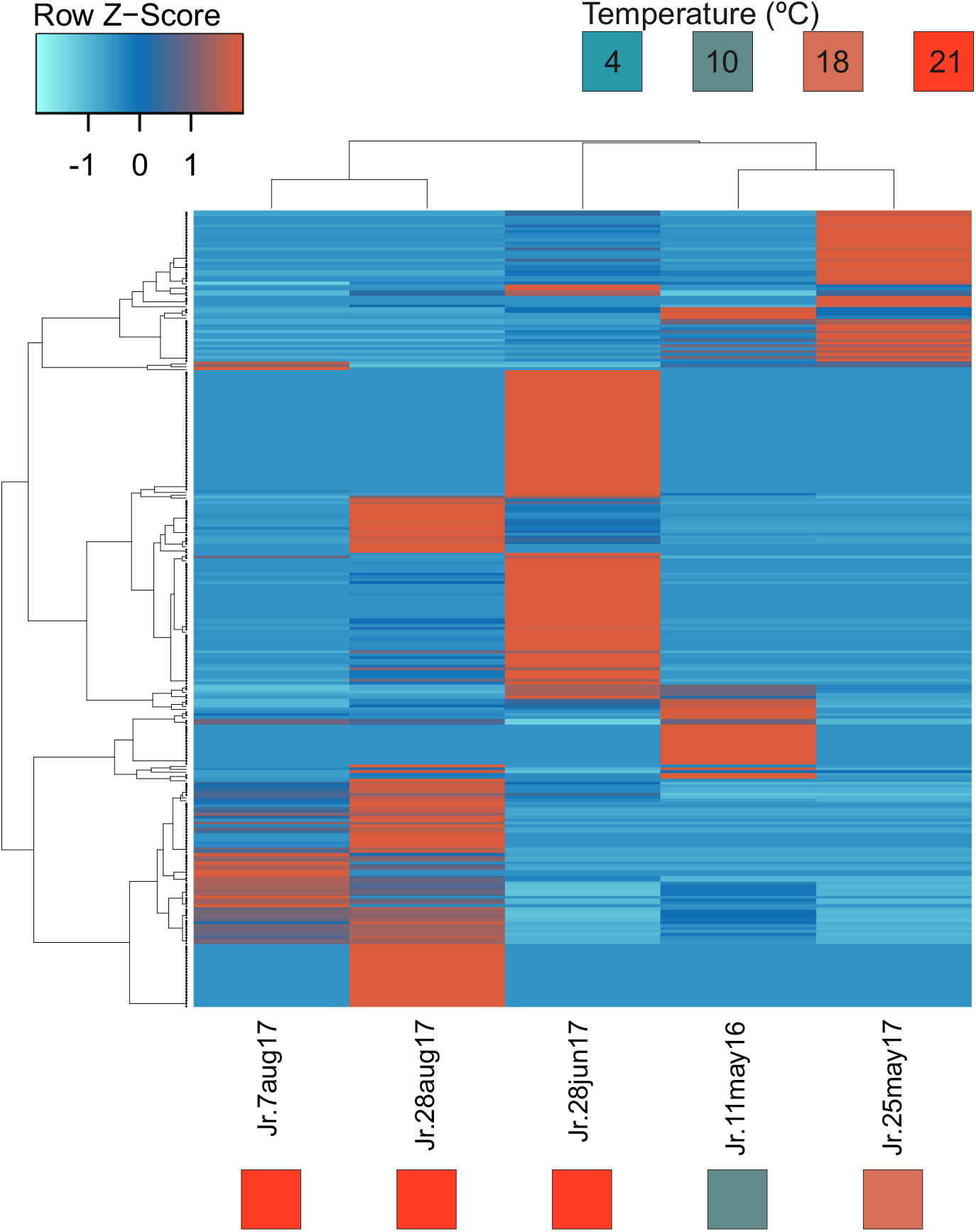
Relative abundance of 279 Jiřická phages in 5 metagenomes (coverage per Gb of metagenome normalized by Z-score). Phages are clustered by sample and abundance by average linkage (with Spearman Rank correlation method). Columns are annotated with the temperature and the sampling date. Temperature color key is shown at top right.

## List of Supplementary Tables

**Supplementary Table S1**: Metagenomic datasets used in this study

**Supplementary Table S2**: Freshwater phage genome information

**Supplementary Table S3**: Phage Genomes encoding ADP-ribosyltransferase toxin (pfam domain PF03496)

**Supplementary Table S4**: Freshwater phage genomes encoding ribosomal proteins

**Supplementary Table S5**: Oxidative Stress Related Pfam Domains

**Supplementary Table S6**: Genome statistics and abundances of 444 actinobacterial bins

**Supplementary Table S7**: TIGRFAMs markers used for phylogenomic analyses

